# CoreProfiler: core genome Multilocus Sequence Typing combining local allele calling and reference nomenclature synchronization

**DOI:** 10.64898/2026.11.26.769085

**Authors:** Alix de Thoisy, Julie Lao, Cléa Siguret, Hugo Lefeuvre, Bérénice Batut, Federica Palma, Philippe Glaser, Claudine Médigue, Fabien Mareuil, Alexis Criscuolo, Sylvain Brisse

## Abstract

Whole-genome sequencing of bacterial isolates enables high-resolution strain genotyping for epidemiological surveillance and outbreak investigations. To convert sequences into comparable genotypes, multilocus sequence typing (MLST) and its more discriminatory extension, known as core genome MLST (cgMLST), record allele identifiers found in genomes at predefined sets of loci, known as schemes. Reference nomenclature platforms provide definitions of alleles and their unique combinations, known as sequence types. These public nomenclatures can be used locally for confidential genotyping. However, standalone cgMLST genotyping tools face challenges such as scalability and the need for a standardized definition of new alleles. Here, we introduce CoreProfiler, a tool for (cg)MLST-based typing of large batches of bacterial genomes using an efficient two-step BLAST approach. Alleles already present in the reference nomenclature are reported with their official identifiers, whereas novel alleles are defined using a hashing approach, ensuring comparability with future reference alleles. CoreProfiler proved to be fast and accurate in both our benchmark and an independent one. CoreProfiler is available as a standalone command-line tool as well as a Galaxy tool and workflow, and is integrated into the ABRomics platform (https://www.abromics.fr) dedicated to research and surveillance on antimicrobial resistance. By combining fast, accurate, and confidential local genotyping with comparability to reference nomenclatures, CoreProfiler offers a flexible solution for standardized yet decentralized genomic surveillance.

**Impact statement:** CoreProfiler addresses a critical bottleneck in genomic epidemiology by enabling fast and accurate local bacterial genotyping, compatible with central nomenclature databases. By implementing an accurate BLAST-based approach for large-scale allele calling, CoreProfiler allows users to process genome assemblies of clinical isolates locally, bypassing the data confidentiality issues associated with submission to central platforms. It also bridges the gap between local genotyping and global references by combining official nomenclature identifiers for known alleles with MD5 hash values for novel ones. This balance ensures that independent public health microbiology actors can investigate outbreaks and compare genotypes among themselves. Integrated into major workflow platforms, CoreProfiler delivers a scalable solution for synchronized global pathogen surveillance.

**Data Summary:** CoreProfiler is implemented in Python (≥ 3.11) and released under the GPL-3 license. Its source code, documentation, and installation instructions are available at https://gitlab.com/ifb-elixirfr/abromics/coreprofiler. It is also accessible as a Galaxy tool at https://usegalaxy.fr/?tool_id=toolshed.g2.bx.psu.edu/repos/iuc/coreprofiler_allele_calling/coreprofiler_allele_calling and as a Galaxy workflow at https://iwc.galaxyproject.org/workflow/cgmlst-bacterial-genome-main.

## 1. Introduction

Defining the precise genetic identity of bacterial isolates that cause clinical infections is crucial for elucidating their sources, transmission patterns, and epidemiological emergence. While whole-genome sequencing is now routinely performed, comparisons of sequence data require standardized methods. Gene-by-gene approaches such as multilocus sequence typing (MLST) [1] and its extension to the core genome (i.e., the set of gene loci present in all genomes within a bacterial taxon; cgMLST) [2, 3] are highly standardized and reproducible: genomes are compared using a fixed scheme, *i.e*., a set of predefined core gene loci (typically, full protein-coding sequences) at which sequence variation is recorded. Allele calling is a fundamental component of (cg)MLST, as it defines the exact nucleotide sequence version of each locus within genome sequences and records the corresponding allele identifier. Furthermore, allele calling implies the discovery and recording of novel alleles. The comparison of the allele profiles of two genomes yields a pairwise allele distance value (i.e., the number of loci at which the two profiles carry different alleles), a key metric in assessing genetic relatedness and a basis for grouping bacterial pathogens into “strains”, as defined for outbreak investigation, surveillance, population biology, and taxonomy.

To enable the comparison of bacterial isolates’ genomes, genotyping schemes and their reference allele sequences are made accessible to the community of investigators using several models. Reference platforms, such as PubMLST [4], BIGSdb-Pasteur [5], and Enterobase [6, 7], maintain public reference nomenclatures of alleles and profiles, together with groupings derived from them, e.g., profile clusters or Life Identification Number (LIN) codes [5]. Allele calling from genomic data is performed by these reference platforms upon genome submission, and curators validate the resulting new allele sequences before they are integrated into the nomenclature. However, this model may in some cases raise concerns regarding privacy, as submission of genome sequence data to these reference platforms is required. To avoid this confidentiality breach issue, it is possible to decentralize the allele calling step by downloading the reference allele sequences (and associated identifiers) from reference nomenclature platforms in order to perform local genotyping. Once retrieved, alleles can then be searched for in local genome sequences. Tools that call alleles locally against reference MLST schemes include *mlst* (https://github.com/tseemann/mlst) and a few others [8], but they are restricted to classical MLST schemes (with 5 to 8 genes, typically 7). Examples of tools developed for genome-scale MLST allele calling include chewBBACA [9, 10] and MiST [11]. However, novel alleles identified locally are generally not compared with the reference platforms. To address this limitation, a third model is implemented in frameworks such as chewie-NS [8], which is compatible with chewBBACA allele calling, and cgMLST.org nomenclature servers, the latter being operated via the commercial tool MBioSEQ Ridom Typer from Bruker (previously seqSphere from RIDOM) [12, 13]. Both chewie-NS and cgMLST.org enable local allele sequences to be shared with a reference repository, which can assign them public identifiers, whilst preserving the confidentiality of the genome data. The nomenclature server cgMLST.org also stores allelic profiles and cluster types, defined at a fixed threshold. A caveat of this model is the lack of centralized quality control of the allele sequences. Finally, the “distributed cgMLST” approach has also been proposed, in which the nomenclature itself is decentralized: alleles are named by the MD5 hash of their nucleotide sequence, removing the need for a reference database [14]. Using this latter strategy, identical novel allele sequences are assigned identical identifiers, regardless of where they are identified. A drawback of distributed cgMLST is the lack of a common nomenclature of strain genotypes.

We developed CoreProfiler to address three needs: (i) computational efficiency, (ii) confidentiality of genome data, and (iii) comparability of locally defined allele sequences with reference platform nomenclatures. CoreProfiler is a standalone tool designed for high-efficiency BLAST-based allele calling against reference allele sequences. It performs a fully local, confidential allele calling using any existing reference nomenclature. CoreProfiler also labels novel alleles by using nucleotide sequence hashing, thus making them comparable internally. Initially named CGprofiler, for Core Genome profiler (https://gitlab.pasteur.fr/BEBP/CGprofiler), the tool was subsequently renamed CoreProfiler and integrated into the ABRomics platform (https://www.abromics.fr), where the source code is maintained (https://gitlab.com/ifb-elixirfr/abromics/coreprofiler).

## 2. Design and implementation

### 2.1 Input

CoreProfiler takes as input one genome assembly and one (cg)MLST scheme. The scheme consists of a set of FASTA-formatted locus files (each containing nucleotide allele sequences with associated identifiers). Schemes can be readily downloaded from the chosen reference nomenclature platform. CoreProfiler requires the NCBI BLAST+ suite (v2.16 or later) [15]. Prior to any analysis, the scheme sequences are formatted into a BLAST database using the BLAST+ tool `makeblastdb`. Although fast (~1 s per 100,000 alleles), this initial step needs to be executed only when the scheme is updated. Therefore, users can create their own BLAST database once and specify its path to CoreProfiler.

### 2.2 Allele calling

The allele calling function is split into two parts, inspired by the strategy implemented within the BIGSdb application [4]: `Autotag` to search for known (reference) allele sequences, and `Scannew` to search for new allele sequences (Figure 1). Both allele calling functions are based on BLASTn searches that can be run using multiple threads. Temporary allele sequences remain comparable despite parallelization, given the hashing strategy described below. All BLASTn searches disable low-complexity masking (option: -dust no) to prevent potential false negatives.

**Figure 1.**
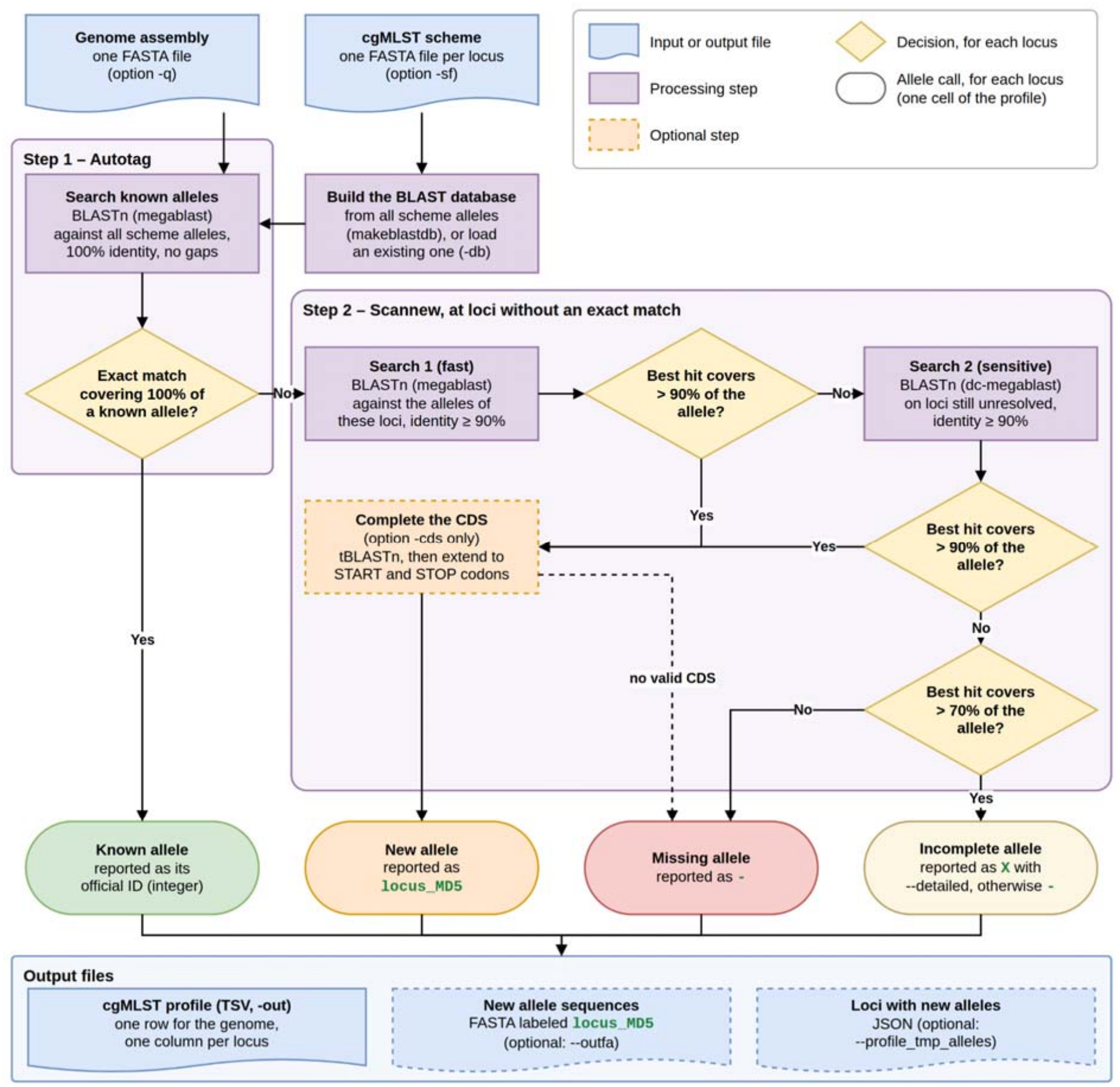
Overview of the cgMLST allele identification workflow design in CoreProfiler. The workflow takes genome assemblies and curated schemes as input. A two-step BLASTn search is then applied: (Step 1) the Autotag function searches for exact allele matches, and (Step 2) the Scannew function identifies potential novel or incomplete alleles at unassigned loci during Step 1. The CoreProfiler output includes a combination of the official allele identifiers (from the reference nomenclature) and the MD5 hashes of the nucleotide sequence for every newly inferred allele.

During the `Autotag` step, CoreProfiler performs a fast BLASTn search of the genome sequence(s) against the scheme allele sequences, using parameters optimized for speed (options: -task megablast, -word_size 31, -perc_identity 100, -ungapped). CoreProfiler next parses the BLASTn results to keep matches with 100% allele coverage. When multiple allele sequences are retained for the same locus (*e.g*., when the reference scheme contains allele sequences that are substrings of one another), CoreProfiler reports only the identifier of the longest matching allele sequence.

The `Scannew` step is subsequently applied only to loci where no known allele sequence was identified during `Autotag`. First, a fast BLASTn search (options: -task megablast, -word_size 28, -perc_identity 90) is performed against all allele sequences of these loci. A new allele is then assigned when the best hit within a locus achieves more than 90% subject allele coverage. Second, the remaining loci (without assigned allele) are searched using more sensitive parameters (options: -task dc-megablast, -word_size 11, -template_length 21, -template_type coding, -perc_identity 90); each of these loci is then characterized based on the subject coverage of its best-scoring hit:

□ coverage > 90%: new allele;
□ coverage > 70% and ≤ 90%: incomplete allele (labeled “X” with the --detailed option);
□ coverage ≤ 70%, or no hit: missing allele (labeled “-”).

For schemes in which every locus is a coding sequence, CoreProfiler also implements the “-cds” option, which applies to all loci for which a new complete allele was found. A new allele that already constitutes a complete coding sequence (i.e., in-frame and bounded by START and STOP codons) is reported unchanged. Otherwise, the best-matching reference allele sequence is translated and used as a query in a tBLASTn search (options: -seg no, -word_size 3) against the contig sequence. Unambiguous hits (i.e., identity > 90% and allele query coverage > 90%) are retained, and the genomic positions of the best one are extended to the flanking START and STOP codons. The new allele is then returned when it is short enough (up to twice the query allele length), and no internal STOP codon occurs; otherwise, a missing allele is reported (labeled “-”).

### 2.3 Identifying temporary alleles as their MD5 for local and external comparability

As CoreProfiler operates locally, it cannot assign official reference nomenclature identifiers to each newly identified allele. Instead, in line with the decentralized cgMLST approach, it provides an MD5 hash of the nucleotide sequence as a temporary identifier. This identifier is a deterministic 128-bit (32-character hexadecimal; https://www.rfc-editor.org/info/rfc1321) fingerprint used both for local comparisons of profiles from genome sequences processed in parallel and for future allele matching. This approach thus guarantees that the same identifier across parallel runs is assigned to each new and identical allele sequence.

### 2.4 Output

CoreProfiler generates a single tab-delimited file containing one column per locus (with the loci listed in alphabetical order), the overall row representing the genome profile. Each cell of the table contains either (i) in case of an exact match, the official allele integer identifier; (ii) for newly inferred alleles, the MD5 hash of the nucleotide sequence; (iii) “X” for incomplete allele sequences (when the --detailed option is enabled); or (iv) “-” for a missing locus. To ease the integration of CoreProfiler results into subsequent analyses, two optional files can also be generated: a FASTA file containing all the newly inferred nucleotide allele sequences, each labeled with its MD5 hash identifier (option --outfa), and a JSON file detailing every locus carrying temporary alleles (option --profiles_tmp_alleles).

### 2.5 Nomenclature synchronization

The temporary identifiers assigned to novel alleles (Section 2.3), together with the sequence and tracking files generated by CoreProfiler (Section 2.4), make locally called profiles synchronizable with the reference nomenclatures over time. The local scheme can be kept up to date using the `db update` command, which retrieves, through authenticated APIs (OAuth1 for PubMLST and BIGSdb-Pasteur), only the loci modified since the last local update.

In standalone usage, profiles that contain temporary (hash-labeled) allele sequences can be resynchronized at any time by re-running CoreProfiler on the corresponding assemblies with the updated scheme: allele sequences integrated into the reference nomenclature in the meantime are then assigned their official identifiers, as observed for the single discordant allele of our validation experiment (see Results).

In platform usage—for example, where genome assemblies are not retained—synchronization can be achieved directly in sequence-hash space: upon each scheme update, the current reference allele sequences of every updated locus may be retrieved and hashed, and any stored temporary identifier whose hash matches a now-official allele can then be replaced by its official integer identifier. This is the approach adopted in the ABRomics platform. Temporary identifiers that have no counterpart yet in the reference nomenclature are left unchanged and remain comparable across independent analyses.

### 2.6 Distribution and integration into Galaxy and the ABRomics platform

CoreProfiler is distributed as an open-source (GNU GPL-3) command-line tool, with a Bioconda recipe to ease its installation and with a bio.tools entry for its findability. To support reproducible, large-scale use, CoreProfiler was wrapped as a Galaxy tool (“coreprofiler_allele_calling”), available on the usegalaxy.fr and usegalaxy.eu servers, and deployable on any Galaxy instance. It is also integrated into a dedicated Galaxy workflow (“cgmlst-bacterial-genome”) maintained by the Intergalactic Workflow Commission (IWC). CoreProfiler, via its Galaxy integration, now serves as the cgMLST engine for the ABRomics platform (https://analysis.abromics.fr), which is used as a French national infrastructure for antimicrobial resistance analysis. In this platform, inferred cgMLST profiles are stored and used for comparative analyses between genome sequences from diverse laboratories, enabling clusters of closely related profiles to be identified. ABRomics computes pairwise dissimilarities between profiles and implements a matching function that will generate alerts for surveillance and outbreak detection purposes. CoreProfiler version 2.0.0 onwards supports authenticated (OAuth1) access to PubMLST and BIGSdb-Pasteur, in addition to EnteroBase and cgMLST.org, to retrieve and update schemes.

## 3. Results

To assess both calling steps (`Autotag` and `Scannew`) under conditions representative of local use, we genotyped using cgMLST, 89 *Klebsiella pneumoniae* genome sequences from a recent genomic surveillance study [16] (Figure 2). CoreProfiler was run locally against the scgMLST629_S scheme from BIGSdb-Pasteur, downloaded on 2024-12-30 (629 loci, 640,296 allele sequences in total, yielding 1,018 sequences per locus on average). The 89 genome sequences were also deposited in BIGSdb-Pasteur to be analyzed using the live nomenclature 19 months later (2026-07-27). Alleles added to the reference BIGSdb nomenclature in the meantime were therefore seen as novel by CoreProfiler.

**Figure 2.**
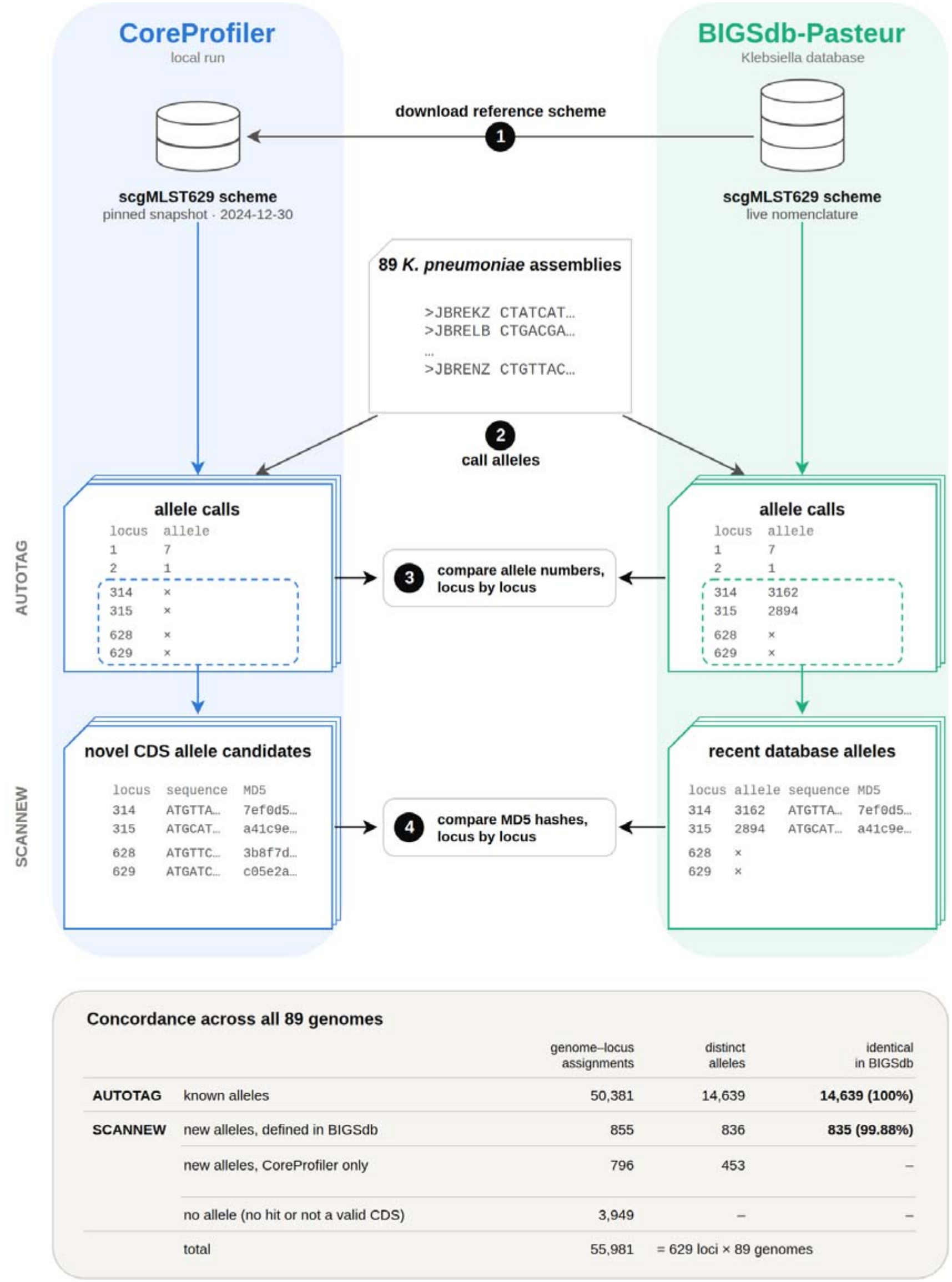
Validation of novel-allele inference (Scannew) against the BIGSdb-Pasteur nomenclature. Eighty-nine *Klebsiella pneumoniae* genome sequences were genotyped by cgMLST using CoreProfiler against a scgMLST629_S scheme snapshot frozen on 2024-12-30 and, independently, using BIGSdb-Pasteur based on the live nomenclature retrieved on 2026-07-27. Locus calls fall into three classes: (i) identifiers returned by both tools, compared directly; (ii) CoreProfiler MD5 hashes against a BIGSdb-Pasteur identifier, compared after hashing the curated sequence; and (iii) novel alleles with no reference identifier, which cannot be assessed. Counts are summed over the 89 genomes. A cross (x) indicates an absence of allele.

The 89 genomes represent 55,981 (=629×89) locus calls. For existing alleles (identified by `Autotag`), CoreProfiler resolved 50,381 of these calls (90.0%) to an existing allele identifier, all identical to those assigned by BIGSdb-Pasteur. The remaining 5,600 (10.0%) were then processed by `Scannew` with the -cds option, yielding a novel allele for 1,651 of them (1,289 distinct sequences, at 446 loci) and no allele (“-”) for the remaining 3,949 (also 446 distinct loci by coincidence, although another set than for the novel alleles). For 836 of the 1,289 novel allele sequences (64.8%, found in 855 genome-locus combinations), BIGSdb-Pasteur had also defined an allele: 76 earlier from other isolates and 760 from these same genomes. All the 836 alleles except one were identical to the allele sequences inferred by CoreProfiler. The only discordance concerned locus ycfS_S, which has two in-frame START codons six bases apart: CoreProfiler delineated a novel allele from its best BLAST hit among all reference alleles of the locus, here starting at the first START codon, whereas BIGSdb-Pasteur delineated it from its closest type allele, which starts at the second START codon. For the other 453 (=1,289–836) novel allele sequences inferred by CoreProfiler (796 genome-locus combinations, at 101 loci), BIGSdb-Pasteur defined no allele, because their length exceeded the length limits set for the corresponding loci (+/-5% of standard sequence length).

In terms of running times, the analysis of the 89 genomes was completed in 13 minutes by CoreProfiler using eight threads (9.0 s per genome, with a maximum memory usage of 6.7 GB) on an Apple M4 Pro (14 cores) with 48 GB RAM. Steps `Autotag` and `Scannew` (using the -cds option) accounted for 17% and 83% (1.5 s and 7.5 s per genome, on average) of the overall running time, respectively. The `Scannew` step runtime increased linearly with the number of loci left unresolved by `Autotag`, which averaged 63 of 629 loci per genome in this dataset.

## 4. Discussion

CoreProfiler was developed to meet the growing need for accurate and efficient cgMLST genotyping of large sets of assembled genome sequences, while combining the needs for data confidentiality and for comparability with reference allele nomenclatures. It was engineered for integration into other tools and genomic surveillance platforms, such as ABRomics.

The accuracy of novel allele definition proved nearly total compared to the reference `Scannew` method from BIGSdb, using a recent real-case *K. pneumoniae* genome sequence dataset: a local scheme 19 months out of date differed from the reference nomenclature at a single novel allele out of 836, and re-synchronization resolved this unique difference, resulting in perfect agreement between the alleles called by CoreProfiler and by the caller of the reference BIGSdb database.

A third-party benchmark of cgMLST accuracy to call known alleles was published recently [11], showing that CoreProfiler matched the correct allele at every locus of every scheme evaluated, including the largest ones (up to ~3,000 loci). CoreProfiler also achieved the highest accuracy among the tools compared, together with the online platform BIGSdb [11].

This benchmark also measured CPU time, wall-clock time, total Input/Output (I/O), and peak memory, with all tools run on four threads against pre-indexed scheme databases. It was found that CoreProfiler (version 1.1.3, which is approximately twice as slow as the version 2.0.0) was more than twice as fast as chewBBACA for allele calling, with about a quarter of its I/O footprint, whilst achieving significantly higher accuracy [11]. The allele calling implemented in chewBBACA failed at short loci, due to its annotation-first strategy. Another tool, MiST, developed more recently based on the minimap2 sequence alignment tool [17], has proved to be around twice faster (than CoreProfiler v1.1.3) while requiring less memory. Besides, CoreProfiler remained more accurate: MiST differed from the reference method for 31 of 118,080 calls (99.97% accuracy), mostly at short *E. coli* loci (45-63 bps), whereas CoreProfiler matched the reference at every locus.

CoreProfiler therefore appears as a promising tool for fast and accurate cgMLST-based genotyping, where profiles must match the reference nomenclature alleles exactly. Its two-step `Scannew` search ensures that the detection of novel alleles is fast, with a second, more sensitive search that should benefit schemes with short allele sequences or highly polymorphic loci.

A limitation of CoreProfiler is that, as for all assembly-based methods (e.g., tools like chewBBACA, BIGSdb, pyMLST [18], or MiST), this tool is sensitive to potential assembly errors in the input contig sequences. Conversely, read-based allele calling tools, e.g., SRST2 [19], ARIBA [20], KMA [21], or stringMLST [22], require as input the raw sequencing reads, which are large files that can be difficult to share. The assembly-based tools also differ importantly in how alleles are called: chewBBACA predicts coding regions in the assembled contig sequences before matching them to the scheme, whereas alignment-based tools, such as BIGSdb, use BLASTn [23] to directly align the reference allele sequences against the contigs. Although CoreProfiler was not specifically designed to run on classical (7 gene-based) MLST schemes or other schemes with loci that are not complete coding sequences, it can be used for these loci without the -cds option.

CoreProfiler is a cgMLST tool that aims at reconciling local allele calling (thus preserving confidentiality of genomic sequences) with comparability with reference nomenclatures hosted and maintained in PubMLST, BIGSdb-Pasteur or EnteroBase. Similar to CoreProfiler, MiST is an efficient and accurate cgMLST tool, which additionally provides the flexibility needed to genotype non-coding sequence loci. However, its accuracy for defining novel alleles (equivalent to `Scannew`) has not been evaluated, to our knowledge. ChewBBACA is an advanced tool to design cgMLST schemes and for allele calling, but its annotation-first strategy creates some incompatibility with alignment-first allele calls [11]. Other cgMLST frameworks such as chewie-NS and cgMLST.org maintain their own nomenclatures, to which users can add new alleles (with cgMLST.org also providing shared cluster types), but these novel alleles cannot be directly matched with reference nomenclature alleles and profiles from PubMLST, BIGSdb-Pasteur and EnteroBase due to different allele calling algorithms (chewie-NS, which relies on chewBBACA) or scheme loci (cgMLST.org). We note that MiST, which was published while this work was in progress, has an allele sequence hashing strategy [11] similar to CoreProfiler, also enabling forward-comparability of novel alleles from reference databases.

In conclusion, CoreProfiler is a practical open-source tool for highly accurate and local cgMLST genotyping that enables retrospective and prospective allele matching with the curated reference nomenclatures, on which strain taxonomic elements such as cgST profiles, hierarchical clusters, and LIN codes are defined [5, 7, 24]. It may thus contribute to resolving the current dilemma between the need to keep sequence data local, for confidentiality reasons, and the need to communicate globally on genotypes defined in public strain nomenclature databases. CoreProfiler could therefore facilitate the identification of emerging strains of public health importance and the build-up of collective knowledge on their epidemiology and evolution.

## Open access

This research was funded, in whole or in part, by Institut Pasteur and by the European Union’s Horizon 2020 research and innovation programme. For the purpose of open access, the authors have applied a CC-BY public copyright license to any Author Manuscript version arising from this submission.

## Author statements

### Author contributions

Conceptualization: A.dT., S.B., A.C.; Software: A.dT., J.L., F.M., B.B., C.S., H.L.; Validation: A.C., F.P.; Funding acquisition: P.G., C.M.; Project administration: S.B., P.G., C.M.; Supervision: S.B., F.M., P.G., C.M.; Writing original draft: A.dT., S.B.; Writing, review and editing: all authors.

## Acknowledgements

We are indebted to Institut Français de Bioinformatique for support in the installation and maintenance of the ABRomics platform, and the Galaxy tools and workflows, in which CoreProfiler was included in the cgMLST workflow.

## References

1. Maiden MC, Bygraves JA, Feil E, Morelli G, Russell JE, et al. Multilocus sequence typing: a portable approach to the identification of clones within populations of pathogenic microorganisms. Proc Natl Acad Sci U A 1998;95:3140–5.

2. Moura A, Criscuolo A, Pouseele H, Maury MM, Leclercq A, et al. Whole genome-based population biology and epidemiological surveillance of Listeria monocytogenes. Nat Microbiol 2016;2:16185.

3. Maiden MC, van Rensburg MJ, Bray JE, Earle SG, Ford SA, et al. MLST revisited: the gene-by-gene approach to bacterial genomics. Nat Rev Microbiol 2013;11:728–36.

4. Jolley KA, Maiden MCJ. BIGSdb: Scalable analysis of bacterial genome variation at the population level. BMC Bioinformatics 2010;11:595.

5. Palma F, Hennart M, Jolley KA, Crestani C, Wyres KL, et al. Life Identification Numbers: A strain nomenclature approach to aid epidemiological surveillance of bacterial pathogens. PLoS Biol 2026;24:e3003781.

6. Achtman M, Zhou Z, Charlesworth J, Baxter L. EnteroBase: hierarchical clustering of 100 000s of bacterial genomes into species/subspecies and populations. Philos Trans R Soc Lond B Biol Sci 2022;377:20210240.

7. Dyer NP, Päuker B, Baxter L, Gupta A, Bunk B, et al. EnteroBase in 2025: exploring the genomic epidemiology of bacterial pathogens. Nucleic Acids Res 2025;53:D757–D762.

8. Page AJ, Alikhan N-F, Carleton HA, Seemann T, Keane JA, et al. Comparison of classical multi-locus sequence typing software for next-generation sequencing data. Microb Genomics 2017;3:e000124.

9. Mamede R, Vila-Cerqueira P, Carriço JA, Ramirez M. chewBBACA 3: lowering the barrier for scalable and detailed whole- and core-genome multilocus sequence typing. Genome Med 2026;18:51.

10. Silva M, Machado MP, Silva DN, Rossi M, Moran-Gilad J, et al. chewBBACA: A complete suite for gene-by-gene schema creation and strain identification. Microb Genomics. Epub ahead of print 15 March 2018. DOI: 10.1099/mgen.0.000166.

11. Bogaerts B, Roosens NHC, Vanneste K. MiST: rapid, accurate and flexible (core-genome) multi-locus sequence typing (MLST) allele calling from draft genomes. BMC Genomics 2025;26:1106.

12. Jünemann S, Sedlazeck FJ, Prior K, Albersmeier A, John U, et al. Updating benchtop sequencing performance comparison. Nat Biotechnol 2013;31:294–296.

13. Kohl TA, Harmsen D, Rothgänger J, Walker T, Diel R, et al. Harmonized Genome Wide Typing of Tubercle Bacilli Using a Web-Based Gene-By-Gene Nomenclature System. EBioMedicine 2018;34:131–138.

14. Zhong L, Zhang M, Sun L, Yang Y, Wang B, et al. Distributed genotyping and clustering of Neisseria strains reveal continual emergence of epidemic meningococcus over a century. Nat Commun 2023;14:7706.

15. Camacho C, Coulouris G, Avagyan V, Ma N, Papadopoulos J, et al. BLAST+: architecture and applications. BMC Bioinformatics 2009;10:421.

16. White RT, Bakker S, Burton M, Dyet K, Elvy J, et al. Decentralized nanopore genomics reveals diverse Klebsiella pneumoniae and no evidence of patient–patient transmission in a New Zealand hospital. Microb Genomics 2026;12:001700.

17. Li H. Minimap2: pairwise alignment for nucleotide sequences. Bioinforma Oxf Engl 2018;34:3094–3100.

18. Biguenet A, Bordy A, Atchon A, Hocquet D, Valot B. Introduction and benchmarking of pyMLST: open-source software for assessing bacterial clonality using core genome MLST. Microb Genomics 2023;9:001126.

19. Inouye M, Dashnow H, Raven L-A, Schultz MB, Pope BJ, et al. SRST2: Rapid genomic surveillance for public health and hospital microbiology labs. Genome Med 2014;6:90.

20. Hunt M, Mather AE, Sánchez-Busó L, Page AJ, Parkhill J, et al. ARIBA: rapid antimicrobial resistance genotyping directly from sequencing reads. Microb Genomics 2017;3:e000131.

21. Clausen PTLC, Hallgren MB, Overballe-Petersen S, Marcelino VR, Hasman H, et al. Assembly-free typing of Nanopore and Illumina data through proximity scoring with KMA. NAR Genomics Bioinforma 2025;7:lqaf116.

22. Gupta A, Jordan IK, Rishishwar L. stringMLST: a fast k-mer based tool for multilocus sequence typing. Bioinformatics 2017;33:119–121.

23. McGinnis S, Madden TL. BLAST: at the core of a powerful and diverse set of sequence analysis tools. Nucleic Acids Res 2004;32:W20–W25.

24. Unitt A, Krisna MA, Parfitt KM, Jolley KA, Maiden MCJ, et al. Neisseria gonorrhoeae LIN codes provide a robust, multi-resolution lineage nomenclature. eLife 2025;14:RP107758.

